# The core herpes simplex-1 fusion complex drives cell-to-cell spreading of pathological Tau

**DOI:** 10.64898/2026.01.21.700770

**Authors:** Stefanie-Elisabeth Heumüller, Mikhail Sushkin, Elisa Sanchez-Sendin, André Hossinger, Dominik Stappert, Nazlican Altinisik, Louisa Seiwert, Lydia Paulsen, Adalbert Krawczyk, Barbara G. Klupp, Thomas C. Mettenleiter, Philip Denner, Harald Prüss, Ina M. Vorberg

**Affiliations:** German Center for Neurodegenerative Diseases (DZNE), Venusberg Campus 1/ 99, 53127 Bonn, Germany; Department of Neurology and Experimental Neurology, Charité - Universitätsmedizin Berlin; German Center for Neurodegenerative Diseases (DZNE), Berlin, Germany; Institute for Virology, University Hospital Essen, University of Duisburg-Essen, Essen, Germany; Department of Infectious Diseases, University Hospital Essen, University of Duisburg-Essen, Essen, Germany; Institute of Molecular Virology and Cell Biology, Friedrich-Loeffler-Institute, Greifswald-Insel Riems, Germany; Department of Neurology, Rheinische Friedrich-Wilhelms-Universität Bonn, 53127 Bonn, Germany

**Keywords:** Prion-like, Tau, Sup35, herpes simplex virus, neurodegeneration

## Abstract

Neurodegenerative diseases such as Alzheimer’s disease (AD) are characterized by the pathological aggregation of the host-encoded protein Tau into amyloid fibrils. Pathologic protein aggregates composed of Tau are able to spread from cell-to-cell, thereby contributing to disease progression. The exact mechanisms of intercellular dissemination remain ill-defined. Mounting evidence links the herpes simplex virus-1 (HSV-1) to the aetiology of AD. HSV-1 is a prevalent neurotropic virus that replicates in epithelial cells at the site of infection, and subsequently establishes lifelong latency in the peripheral and central nervous systems. In the quiescent latent state, AD brains show elevated expression of structural viral proteins in the absence of virus production. We hypothesized that HSV-1 glycoproteins involved in viral attachment and fusion facilitate the intercellular spreading of proteopathic seeds. Using cellular models, we demonstrate that expression of the HSV-1 core fusion complex, essential for viral entry and direct cell-to-cell transmission, is sufficient to promote aggregate dissemination between cells. Moreover, anti-HSV-1 antibodies present in the cerebrospinal fluid of patients with HSV-1 encephalitis efficiently neutralize viral infection, block viral cell-to-cell transmission, and inhibit propagation of pathological Tau. Thus, latent HSV-1 infection of the brain could contribute to neurodegenerative disease progression, and antiviral antibodies may represent a potential therapeutic strategy.

## Introduction

Alzheimer’s disease (AD) is the most common cause of dementia, affecting more than 55 million people worldwide. The accumulation of amyloid-β and phosphorylated Tau are key events that ultimately cause neuronal death and cognitive decline. Tau deposition and pathological hyperphosphorylation are also present in around twenty other neurodegenerative diseases, collectively termed tauopathies ^1^. Tau is a microtubule-binding protein that is involved in stabilising the cytoskeleton. During disease progression, hyperphosphorylated Tau dissociates from microtubules and rearranges into highly ordered amyloid fibrils, which are deposited in neurons as neurofibrillary tangles. Misfolded Tau can spread to unaffected cells, thereby inducing further Tau misfolding and formation of insoluble aggregates ^2^.

Despite decades of research, the cause of sporadic neurodegenerative diseases remains unknown. It has long been suspected that pathogens such as viruses could play a role in the pathophysiology of dementia ^3^. There is increasing evidence that herpes simplex virus type 1 (HSV-1) infection is a risk factor for developing AD later in life. HSV-1 is a highly prevalent alpha herpesvirus that is usually acquired during early childhood, affecting over 3.7 billion people worldwide ^4^. HSV-1 typically causes mild, recurring cold sores, but can also present with severe manifestations, including herpes keratitis, encephalitis, and neonatal herpes. HSV-1 initially replicates in the epithelial cells of the mucosa and skin, before moving to the sensory neurons of the trigeminal ganglia and the neurons of the central nervous system (CNS), where it establishes lifelong latency ^5–7^. Periodic reactivation in response to stress stimuli produces infectious virions that are transported anterogradely through the axons to the mucosal epithelium, causing recurrent skin and mucosal lesions ^8^. Reactivation might also occur asymptomatically ^9^.

Several lines of evidence suggest that HSV-1 infection contributes to dementia. A meta-analysis demonstrated an association of HSV infections with an increased incidence of dementia ^10^. Further, a recent analysis of DNA sequence data showed that a higher genomic HSV-1 load in brain tissue was associated with a higher risk of developing AD ^11^. Individuals with untreated HSV-1 infection have a twofold increased risk of developing cognitive dysfunction or dementia ^12^. Reactivation of latent HSV-1 also appears to be more frequent in patients suffering from AD ^13^. In line with this, patients with mild cognitive decline and dementia have higher anti-HSV IgG titers, but not IgM titers, arguing for HSV reactivation ^10^. Of note, also conflicting data exist regarding the role of HSV-1 in the development of dementia ^14, 15^. Experimentally, HSV-1 infection leads to an increase in amyloid-β production and Tau hyperphosphorylation (p-Tau) and Tau oligomerization in cells, brain organoids and/or animal models ^16–19^. This led to the assumption that amyloid-ß and p-Tau may have antimicrobial properties ^19–22^. In line with this, antiviral drugs can lower amyloid pathology in mice ^23^.

Herpesviruses have evolved two mechanisms to spread from infected to uninfected cells: infection by cell-free virions, or transmission directly via cell-cell contact sites ^24^. HSV-1 entry requires attachment to cognate receptors on the host cell and the subsequent release of the capsid into the cytosol. The fusion process is coordinated by a complex machinery composed of viral glycoproteins within the virion envelope, which fuses the viral membrane with that of the host cell. The core fusion machinery, consisting of four glycoproteins, is sufficient for viral entry ^25^. When the viral protein gD binds to its receptor, it undergoes a conformational switch that activates the gH/gL heterodimer. The main fusogen, gB, then triggers the fusion of the viral membrane with the plasma or endosomal membrane. The capsid is transported along microtubules to the nuclear pores through which the viral DNA is injected and DNA replication begins. During the productive, lytic viral replication, viral capsids assemble in the nucleus and are released following a complex process involving membrane envelopment at the inner nuclear membrane, de-envelopment at the outer nuclear membrane, and re-envelopment in the trans-Golgi network with the final, glycoprotein-containing envelope ^26^. Subsequent transmission of virus from epithelial cells to sensory neurons depends on direct cell-cell contact ^27^ and is mediated by viral glycoprotein-receptor interactions ^28^. Herpesvirus establishes lifelong latency in the trigeminal ganglia and CNS ^29^. Infectious virions produced during periodic reactivation travel down the axons for reinfection of mucosal epithelial cells. During latency, transcription of lytic phase genes and expression of lytic proteins is generally silenced ^30^. However, latency is not absolute, and expression of lytic genes can resume in the absence of productive reactivation ^31^. Recently, abundant expression of lytic viral structural proteins, such as gB in human brain in the absence of active virus production has been reported ^22^.

We have previously shown that viral glycoproteins associated with exogenous and endogenous enveloped viruses accelerate the spreading of proteopathic seeds by enabling efficient intercellular membrane contact and subsequent protein aggregate transmission. Likewise, viral glycoproteins sorted onto extracellular vesicles (EVs) loaded with proteopathic seeds facilitate their binding to recipient cells and subsequent endosomal escape of cargo into the cytosol ^32, 33^. Here, we tested the hypothesis that expression of the herpesviral core fusion complex-glycoproteins observed during latency is sufficient to drive the intercellular dissemination of proteopathic seeds. We use two cell-based assays to monitor aggregate spreading between cultured cells. In the NM model, we study the mammalian expression of the *Saccharomyces cerevisiae* prion domain NM of Sup35, a translation termination factor that can adopt an amyloid state. The intrinsically disordered, glutamine/asparagine-rich NM domain lacks mammalian homology but resembles proteins aggregating in neurodegenerative diseases (e.g., TDP-43, FUS) ^34^. In the mammalian cytosol, NM remains soluble until seeded by recombinant NM fibrils ^35^, after which aggregates spread from cell-to-cell via direct contact or EVs ^36^. This model enables general studies of prion-like protein transmission. Our second cell-based assay investigates Tau aggregate dissemination. Donor cells express the aggregation-prone Tau repeat domain with P301L/V337M mutations fused to GFP. Stable Tau aggregation in the donor cell line was induced using AD brain homogenate, and Tau seeds spread to recipient cells by contact or EVs ^33^.

We here show that the core HSV-1 fusion complex is sufficient to drive intercellular dissemination of proteopathic seeds by direct cell-cell contact. HSV-1 fusion complex-mediated cell-cell contacts result in efficient transmission of protein aggregates, either through cell-cell fusion or through cell-cell contact with recipient cells. These mechanisms are receptor-dependent, as aggregate induction is prevented in cocultures with recipient cells that are refractory to HSV-1 entry. Importantly, naturally elicited monoclonal anti-HSV-1 antibodies derived from the cerebrospinal fluid (CSF) of patients with acute HSV-1 encephalitis exhibit potent antiviral neutralizing activity and protect against virus-induced cell-to-cell transmission of pathological Tau, opening up new avenues for potential therapeutic intervention in neurodegeneration.

## Results

### The minimal HSV-1 fusion complex supports the intercellular dissemination of model proteopathic seeds

We have previously shown that viral fusion proteins derived from vesicular stomatitis virus^32^, SARS-CoV-2 ^32^, endogenous and exogenous murine leukemia virus (MLV) or human endogenous retroviruses ^33^ strongly increase the intercellular spreading of proteopathic seeds when fusion proteins are expressed by donor cells with proteopathic seeds. Here, we set out to test if the HSV-1 fusion complex can also drive the spreading of proteopathic seeds between cells. The core HSV-1 fusion complex consists of the four viral glycoproteins gD, gH/gL and gB, which act in concert to mediate receptor attachment, fusion with cellular membranes and uptake ^25^. To test if expression of these glycoproteins is sufficient to increase dissemination of diverse cytosolic protein aggregates between donor and recipient cells, we made use of our established HEK cell coculture models persistently propagating HA epitope-tagged NM aggregates (HEK NM-HA^agg^) (**Supplementary Fig. S1a**). The advantage of these cell culture systems is that nearly all cells of the donor population produce protein aggregates, which is a prerequisite for sensitive detection of intercellular protein aggregate induction.

Usually, coculture of donor cells with recipient HEK cells stably expressing soluble NM-GFP (HEK NM-GFP^sol^) (**Supplementary Fig. S1a**) results in induction of NM-GFP aggregates in few recipient cells ^32^. HEK cells express HSV-1 entry receptors and are susceptible to HSV-1 infection ^37^. We tested if expression of individual components or the minimal fusion complex was sufficient to increase aggregate induction in recipient cells. HEK NM-HA^agg^ cells were mock-transfected or transfected with individual HSV-1 plasmids or a combination thereof and donor cells were subsequently cocultured with recipient HEK NM-GFP^sol^ cells (**Fig. 1a**). Western blot analysis confirmed that all proteins were expressed by donor cells. For expression of gL, a construct coding for a V5-tagged protein was used, since appropriate antibodies were not available (**Fig. 1b**). As expected, gB, gD, gL and gH were expressed on or near the cell surface of transfected cells (**Fig. 1c**) ^38^. Coculture of transfected donors with recipient cells demonstrated a drastic increase in aggregate induction in recipient cells when all four plasmids coding for the HSV-1 fusion machinery were cotransfected (**Fig. 1d, e**). This result was also replicated when donor cells were transfected with constructs coding for untagged proteins (**Supplementary Fig. S2a-c**). Interestingly, expression of individual HSV-1 glycoproteins alone caused a slight but significant increase in aggregate induction in recipients (**Fig. 1e**). NM-HA protein levels were unaffected by transfection (**Supplementary Fig. S2d, e**). While syncytia formation was observed in cocultures due to the fusogenic activity of the fusion machinery ^39^, aggregate induction was also observed in individual cells (**Supplementary Fig. S2f**). This suggests that the fusion complex also accelerated intercellular aggregate induction independent of cell fusion events. No aggregate induction in recipients was observed when recipients were cocultured with transfected donor cells expressing soluble NM-HA, confirming that aggregate induction was not induced spontaneously (**Supplementary Fig. S2g, h**).

**Figure 1.**
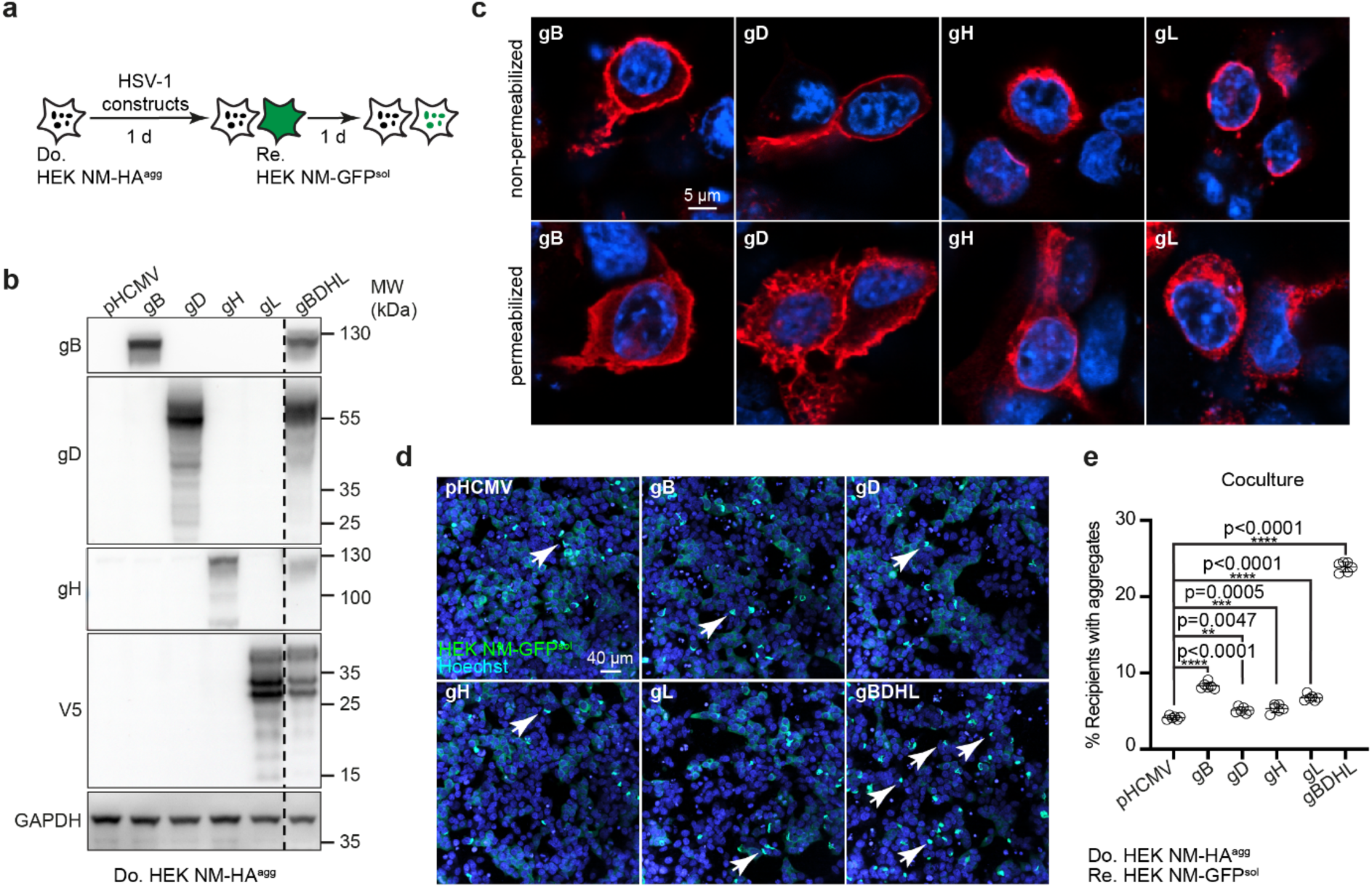
Expression of the HSV-1 fusion machinery in donor cells increases intercellular NM aggregate induction. **a.** Experimental workflow. HEK NM-HA^agg^ donor cells were transfected with individual plasmids coding for gB, gD, gH, gL or a combination of the four plasmids (hereafter termed gBDHL for simplicity) and/or empty vector pHCMV. Donors were subsequently cocultured with recipient HEK NM-GFP^sol^ cells and monitored for aggregate induction. **b.** Western blot analysis of cell lysates from donor cells expressing HSV-1 proteins individually or in combination. gB, gD and gH were detected by specific antibodies, gL was detected using antibodies against fused C-terminal V5-tag. Cell lysates were loaded on two different gels for GAPDH detection on different blots. **c.** Cellular expression of HSV-1 proteins in non-permeabilized and permeabilized donor HEK NM-HA^agg^ cells. Cells were fixed 24 h post transfection. HSV-1 proteins were detected with antibodies used as in (b). Nuclei were stained with Hoechst. **d.** Coculture of donor and recipient cells. Donor cells were not stained in this experiment. Arrowheads depict aggregated NM-GFP. Nuclei were stained with Hoechst. **e.** Quantitative analysis of the percentage of recipient cells with induced aggregates upon coculture. All data are shown as the means ± SD from six replicate cell cultures. Three independent experiments were carried out with similar results. P-values calculated by one-way ANOVA with Dunnett’s multiple comparisons test.

### The minimal HSV-1 fusion complex accelerates the prion-like spreading of Tau misfolding

As a neurodegenerative disease-related cell system, we used our HEK cell model which is based on the stable expression of a soluble four-repeat domain variant of human Tau harbouring P301L/V337M mutations (HEK Tau-GFP^sol^). A donor clone that propagates Tau-GFP aggregates had been established by exposing the cells to AD brain homogenate and subsequent limiting-dilution cloning (HEK Tau-GFP^AD^) ^32^ (**Supplementary Fig. S1b**). Coculture of HEK Tau-GFP^AD^ cells with HEK cells expressing soluble Tau-FusionRed (Tau-FR^sol^) results in few recipients with Tau-FR aggregates ^32^. Expression of the HSV-1 fusion complex in HEK Tau-GFP^AD^ cells did not influence total levels of Tau-GFP, but resulted in an increase in Tau aggregate induction in recipient HEK Tau-FR^sol^ cells (**Fig. 2a-f**). In line with these results, the HSV-1 fusion complex expression in HEK Tau-GFP^AD^ donors induced some syncytia formation (**Supplementary Fig. S2i**). As cell fusion results in cytoplasmic mixing, we quantified cells exhibiting protein aggregates that stained double positive for red and green, as this indicates a fusion event (**Fig. 2e**). We considered cells exhibiting only red Tau aggregates as cells that had received Tau seeds independent of cell fusion (**Fig. 2e**). Approximately half of induced red aggregates were also positive for green, suggesting that both, cell-cell fusion events and cell fusion-independent induction of protein aggregation, occurred (**Fig. 2g**). No Tau aggregate induction was observed when cells expressing soluble Tau-GFP were used as donors (**Supplementary Fig. S2j, k**). Experiments were repeated using primary human astrocytes expressing soluble Tau-FR as recipient cells. Again, coculture with donors expressing the core HSV-1 fusion complex resulted in a strong induction of Tau-FR aggregates in recipients (**Fig. 2h-j**). We conclude that expression of the core HSV-1 fusion machinery by cells harbouring proteopathic seeds is sufficient to induce protein misfolding in cocultured bystanders.

**Figure 2.**
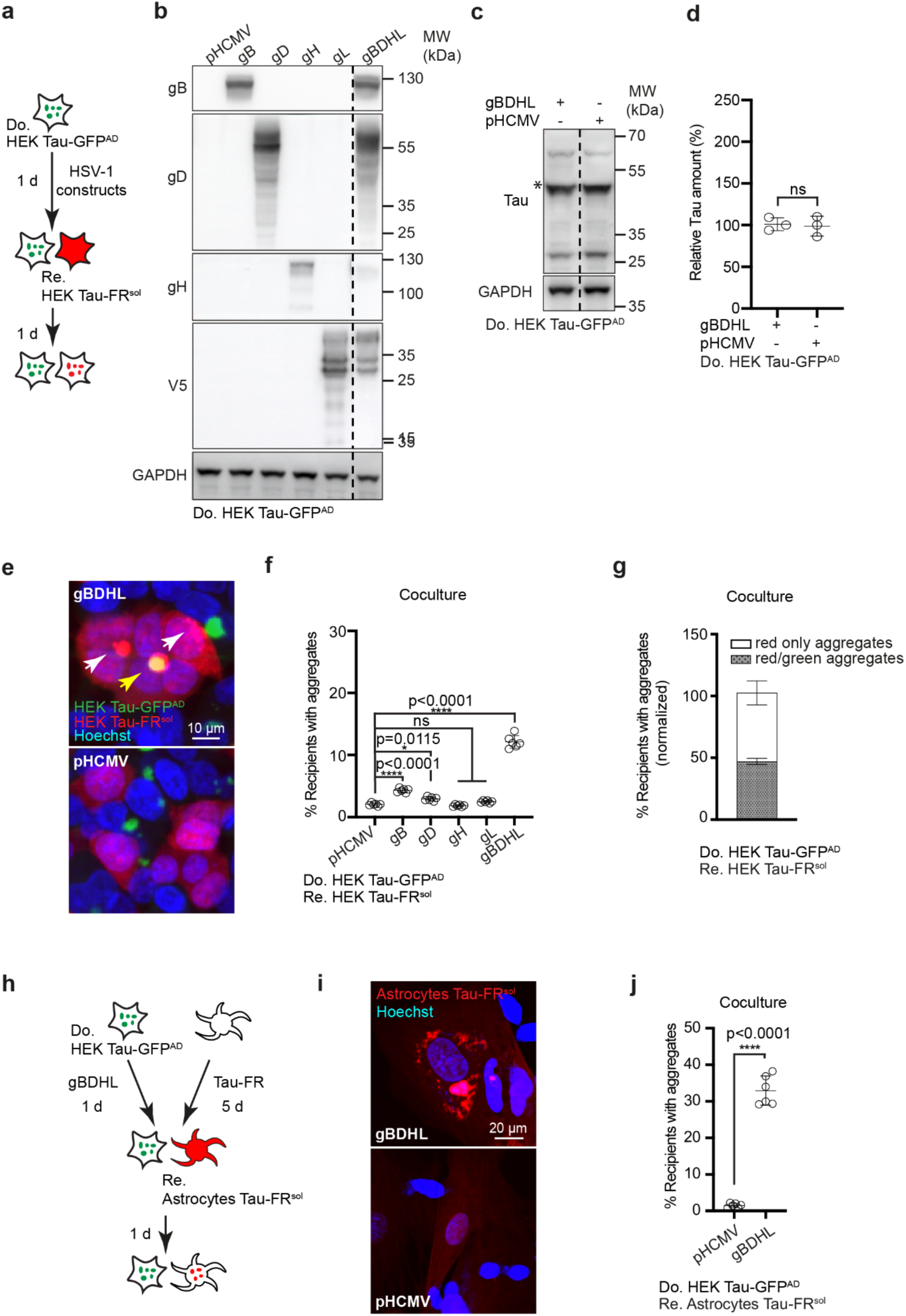
Increased Tau aggregate induction upon expression of the HSV-1 fusion machinery by donor cells. **a.** Experimental workflow. HEK Tau-GFP^AD^ were transfected with plasmids coding for the HSV-1 fusion machinery, and cells were subsequently cocultured with recipient HEK cells expressing Tau-FR^sol^. **b.** Western blot analysis of lysates from the donor cells transfected with plasmids coding for HSV-1 proteins. gB, gD and gH were detected by specific antibodies, gL was detected using antibodies against the C-terminal V5-tag. Samples were loaded on two gels for GAPDH detection on different blots. **c.** Western blot analysis of HEK Tau-GFP^AD^ donor cells transiently transfected with HSV-1 fusion machinery or pHCMV (mock) control. Anti-GFP antibodies were used for Tau-GFP detection. The expected size is indicated by a star. **d.** Quantification of Tau expression in HEK Tau-GFP^AD^ donor cells. All antibody-positive bands were included for the quantification. Mean expression levels in HEK donor cells transfected with mock control were set to 100%. **e.** Coculture of donor and recipient cells. White arrows show examples of ‘red only’ aggregate while the yellow arrow demonstrates a case of double-color ‘red/green aggregate’. **f.** Quantitative analysis of the percentage of recipient cells with induced aggregates. **g.** Quantification of recipients with either red or multinucleated recipients with red and green protein aggregates. **h.** Donor HEK Tau-GFP^AD^ (mock-transfected or transfected with plasmids coding for gBDHL) were cocultured with human primary astrocytes expressing soluble Tau-FR (Astrocytes Tau-FR^sol^) for 1 d. **i.** Confocal images of cocultured human primary astrocytes with induced Tau-FR aggregates when cocultured with gBDHL-expressing donors, but not with pHCMV-transfected donors. **j.** Quantitative analysis of the percentage of astrocytes with Tau-FR aggregates. All data are shown as the means ± SD from six (f, g, j) or three (d) replicate cell cultures. Three (a) or two (h) independent experiments were carried out with similar results. P-values calculated by one-way ANOVA with Dunnett’s multiple comparisons (f) or Student’s t-test (d, j). ns = non-significant.

### HSV-1 fusion machinery-mediated aggregate induction engages viral entry receptors

The foregoing experiments suggested that the increase in intercellular protein aggregate induction is due to efficient contact between donor and recipient membranes mediated by the viral fusion machinery and its entry receptors. We tested this by performing coculture experiments with our NM model using CHO-K1 cells that lack HSV-1 entry receptors as recipients (**Fig. 3a; Supplementary Fig. S3a**) ^40^. Poor NM-GFP aggregate induction was observed when HEK NM-HA^agg^ donors were cocultured with CHO-K1 cells engineered to stably express soluble NM-GFP (**Fig. 3b**). The inefficient aggregate induction was not due to a general resistance of CHO-K1 cells to NM-GFP aggregate induction by exogenous seeds, as exposure of these recipient cells to either recombinant NM fibrils (**Fig. 3c, d; Supplementary Fig. S3b**) or coculture of CHO-K1 NM-GFP^sol^ cells with HEK NM-HA^agg^ cells expressing viral glycoprotein VSV-G (**Fig. 3e, f**) efficiently induced aggregates. Expression of HSV-1 entry receptors HVEM or Nectin-1 ^41^ by recipients strongly increased aggregate induction (**Fig. 3g-i; Supplementary Fig. S3c, d**), confirming that the intercellular spreading of aggregates mediated by the HSV-1 fusion machinery requires expression of viral entry receptors.

**Figure 3.**
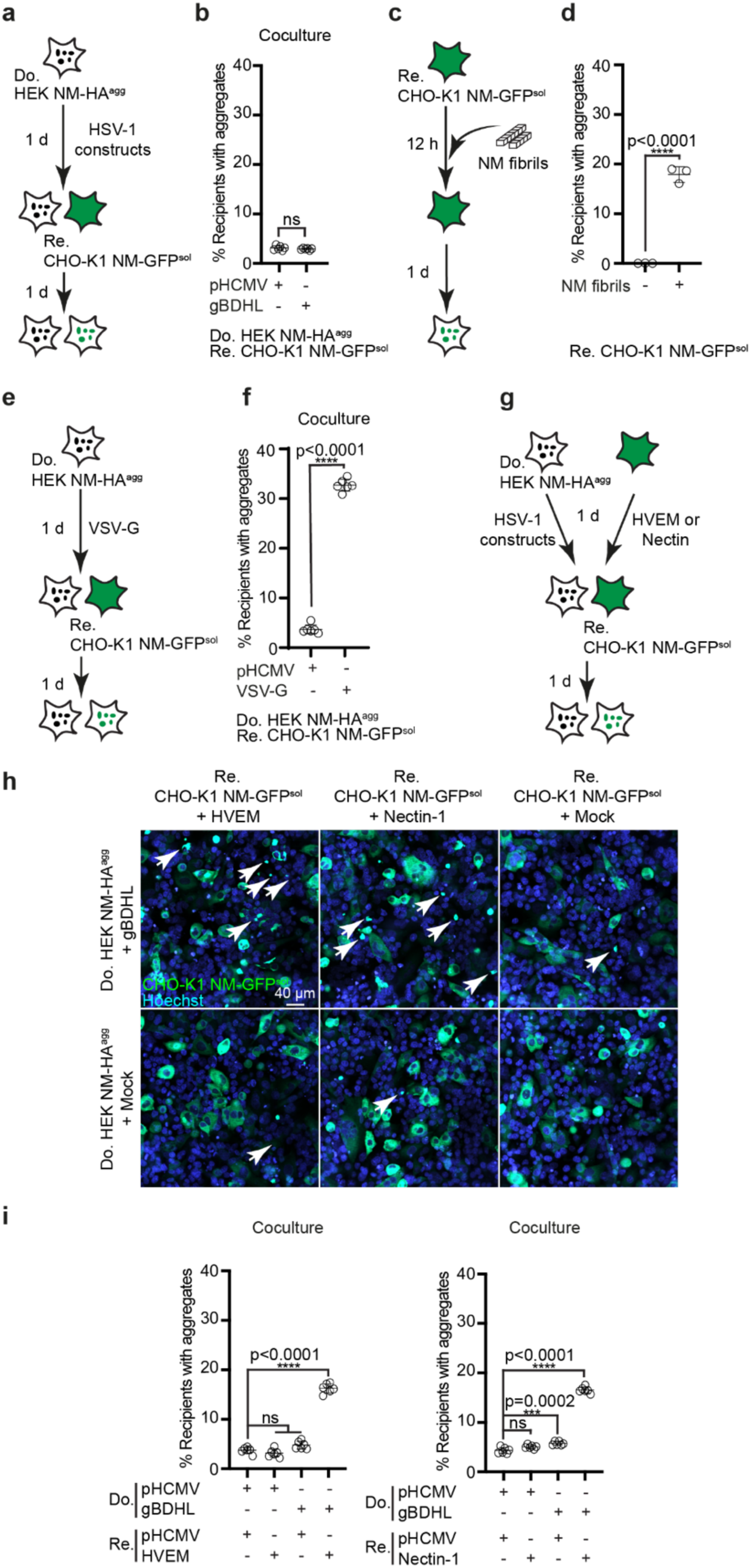
Intercellular proteopathic seed spreading depends on the interaction of the HSV-1 fusion machinery with HSV-1 entry receptors. **a.** HEK NM-HA^agg^ donor cells were transfected with plasmids coding for the HSV-1 fusion machinery and cocultured with HSV-1 resistant CHO-K1 cells stably expressing soluble NM-GFP (CHO-K1 NM-GFP^sol^). Recipients were assessed for NM-GFP aggregate induction. **b.** Percentage of recipient CHO-K1 cells with induced aggregates upon coculture with donors transfected with plasmids coding for the HSV-1 fusion machinery or pHCMV plasmid. **c.** Schematic representation of inducing aggregation in CHO-K1 NM-GFP^sol^ using recombinant NM fibrils. Cells were exposed to 25 µM recombinant NM fibrils (monomer equivalent) and assessed for NM-GFP aggregates the next day. **d**. Successful NM-GFP aggregate induction in CHO-K1 NM-GFP^sol^ cells. Quantification of recipients with NM-GFP^agg^ 24 h post NM fibril exposure. **e.** HEK NM-HA^agg^ cells transfected with VSV-G or pHCMV control plasmid were cocultured with CHO-K1 NM-GFP^sol^ recipient cells. **f.** Quantitative analysis of percentage of recipients with induced aggregates. **g.** Experimental workflow. Donor HEK NM-HA^agg^ cells expressing the HSV-1 fusion machinery were cocultured with recipient cells transiently transfected with plasmids coding for HSV-1 receptors HVEM, Nectin-1 or empty control vector pHCMV. **h.** Coculture of donor and recipient cells. Arrowheads depict aggregates. **i.** Quantitative analysis of the percentage of recipient cells with induced aggregates. All data are shown as the means ± SD from six (b, d, f, i) replicate cell cultures. Three (b, d, f, i) independent experiments were carried out with similar results. P-values calculated by Student’s t-test (b, d, f) or one-way ANOVA with Dunnett’s multiple comparison test (i). ns = non-significant.

### Minimal transmission of proteopathic seeds by conditioned medium and extracellular vesicles

Proteopathic seeds, including pathological Tau and NM aggregates, can be transmitted by EVs to recipient cells, where they act as seeds that induce the aggregation of soluble homotypic protein ^32^. While HEK cells secrete both protein aggregates packaged into EVs, the EV’s ability to induce protein aggregation in recipient cells is usually low, likely due to insufficient uptake and/or poor endosomal escape of cargo ^32^. Both processes can be increased by decoration of donor-derived EVs with viral glycoprotein VSV-G, resulting in efficient aggregate induction in recipient cells ^32^. We tested if the HSV-1 fusion complex would also be secreted in association with EVs (**Fig. 4a, b**). Western blot analysis demonstrated that all four HSV-1 proteins and Tau were present in the EV fraction (**Fig. 4b**). Exposure of recipient cells to these EVs did not result in increased Tau aggregate induction in recipients. Likewise, conditioned medium derived from donor cells expressing the HSV-1 fusion complex only poorly triggered Tau aggregation (**Fig. 4c**). The low Tau aggregate induction by EVs or conditioned medium was not due to diminished secretion of EVs, as donors transfected with the HSV-1 plasmids produced higher levels of EVs secreted into the medium (**Fig. 4d**). We conclude that secretion of proteopathic seeds by donor cells and subsequent uptake by recipients likely does not play a major role in intercellular protein aggregate dissemination mediated by the HSV-1 fusion complex.

**Figure 4.**
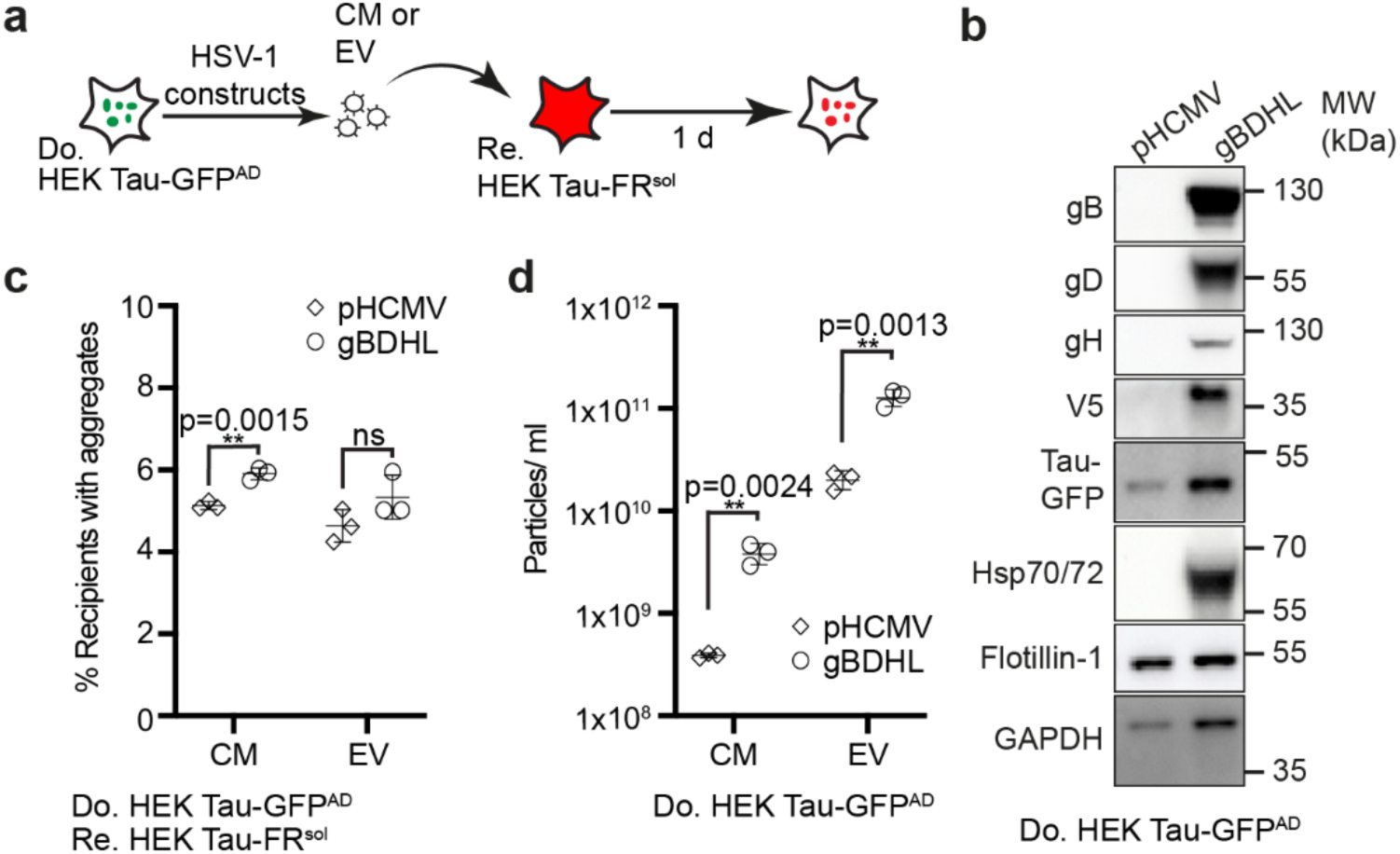
Minor Tau aggregate induction in recipients after incubation with conditioned media and extracellular vesicles collected from HSV-1 fusion complex-transfected donor cells. **a.** Experimental workflow. Donor HEK Tau-GFP^AD^ cells were transfected with pHCMV or with all HSV-1 plasmids coding for gB, gD, gH, and gL. 72 h later, HEK NM-GFP recipient cells were exposed to conditioned medium (CM) or EVs isolated from conditioned medium of HEK Tau-GFP^AD^ donor cells. Tau-FR aggregate induction was determined 24 h post exposure. **b.** Western blot of EV lysates. Equal volumes of EV samples (not adjusted to same particle numbers) were loaded. Flotillin-1 and GAPDH were detected on one Western blot, the other proteins were detected on an additional blot. **c.** Recipient cells were analysed for the percentage of cells with induced Tau-FR aggregates. **d.** Particle numbers in CM and EV samples from HSV-1 fusion complex or pHCMV-transfected donors determined 3 d after transfection. All data are shown as the means ± SD from three (c, d) replicate cell cultures. Three (c, d) independent experiments were carried out with similar results. P-values calculated by Student’s t-test (c, d). ns = non-significant.

### Anti-HSV-1 antibodies block the intercellular dissemination of pathologic Tau

Viral glycoprotein-mediated intercellular transfer of proteopathic seeds can be efficiently blocked by inhibiting the interaction of viral ligands with their cognate cell surface receptors ^33^. Human CSF has been shown to contain immunoglobulins capable of neutralizing HSV-1 virions ^42^. We tested if anti-HSV-1 monoclonal antibodies derived from human CSF could potentially also block intercellular spreading of Tau misfolding. Intrathecal B cells from two patients suffering from acute HSV-1 encephalitis were isolated, single cell mRNA was extracted, and recombinant antibodies preserving epitope specificity were engineered. Screening of cloned antibodies on HSV-1-infected Vero E6 cells (**Fig. 5a**) identified 18 monoclonal antibodies that reacted with HSV-1-infected cells but not with uninfected control cells (**Fig. 5a**; **Supplementary Fig. S4**). Further testing for HSV-1 antigen targets revealed five antibody epitopes on gB, while five antibodies detected epitopes on gD (**Fig. 5b**; **Supplementary Fig. S5**). The antigens for the eight remaining anti-HSV-1 antibodies are unknown.

**Figure 5.**
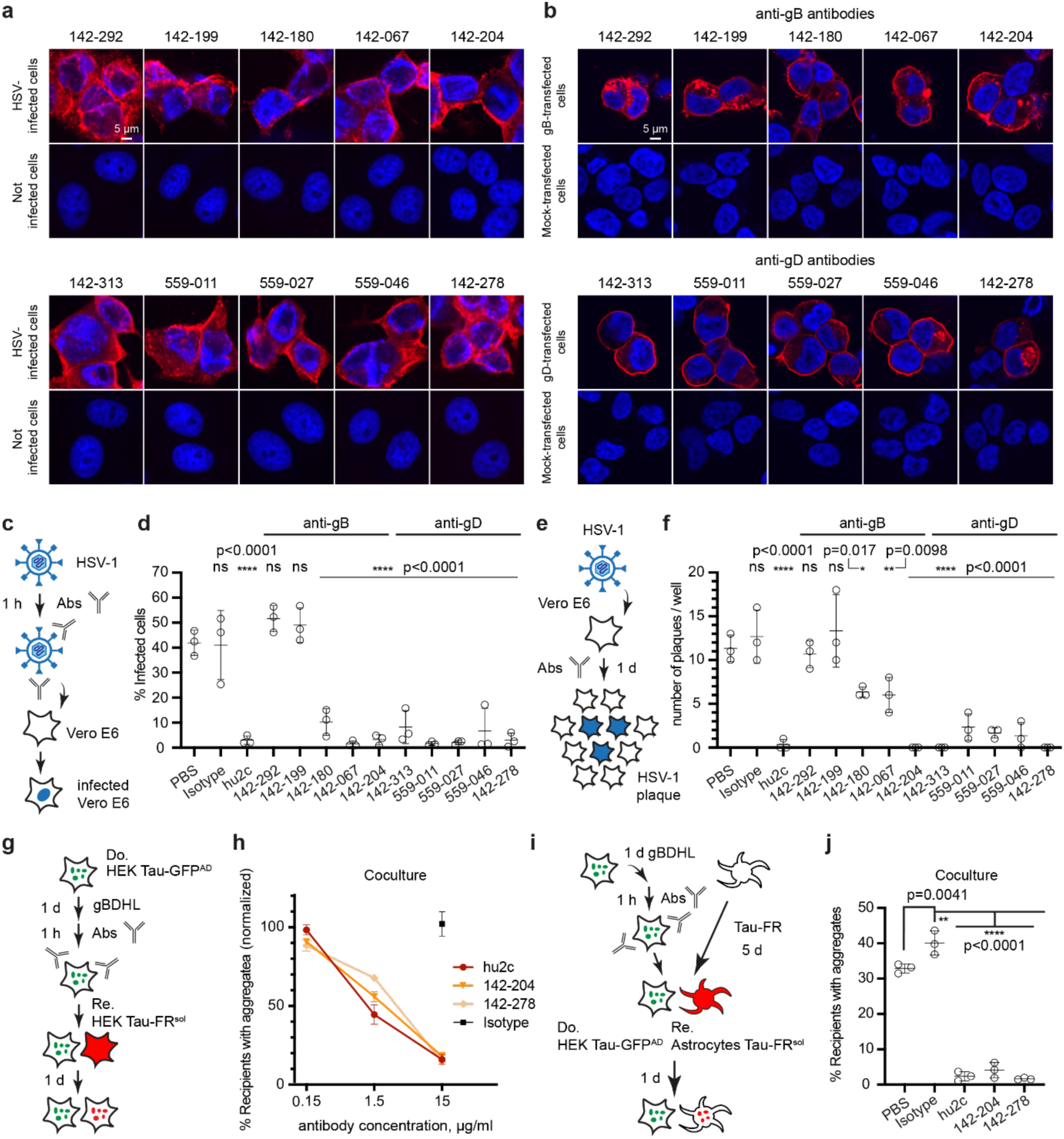
Human CSF-derived monoclonal antibodies against HSV-1 gB and gD possess potent antiviral activity and block the prion-like spreading of proteopathic Tau. **a.** Characterization of recombinant anti-HSV-1 antibodies cloned from CSF-resident B cells of encephalitis patients. B cells from two patients suffering from acute HSV-1 encephalitis were isolated from CSF. Antibodies were subsequently expressed recombinantly in HEK cells. Shown are ten antibodies that bind HSV-1 infected Vero E6 cells but not uninfected control cells. **b.** Characterization of antigens. HEK293T WT cells were transfected with plasmids coding for gB, gD or control vector pHCMV. **c.** Pre-entry inhibition by neutralizing anti-HSV-1 antibodies: HSV-1 was pre-incubated with 15 μg/ml recombinant anti-HSV-1 antibodies, isotype control antibodies, or PBS. Virus was subsequently added to permissive Vero E6 cells. Viral entry was assessed 4 h post infection by staining cells with anti-ICP4 antibodies. **d.** Quantitative analysis of viral infection in the presence of anti-HSV-1 antibodies, determined by ICP4 expression. **e.** Post-entry inhibition of viral infection: Vero E6 cells were first infected with HSV-1 and then incubated with 75 μg/ml anti-HSV-1 antibodies in medium containing 0.6% cellulose. After 24 h incubation with antibodies, the number of HSV-1 plaques per well was counted. **f.** Quantitative analysis of the number of HSV-1 plaques per well. **g.** Coculture of transfected HEK Tau-GFP^AD^ donor cells, preincubated with different concentrations of anti-HSV-1 antibodies, and recipient HEK Tau-FR^sol^ cells. **h.** Normalized percentage of cells with red Tau aggregates following coculture. The percentage of cells (%) with red aggregates incubated without antibodies was set to 100%. **i.** Coculture of transfected HEK Tau-GFP^AD^ donor cells preincubated with anti-HSV-1 antibodies and recipient human primary astrocytes expressing soluble Tau-FR (Astrocytes Tau-FR^sol^). **j.** Percentage of cells with red Tau aggregates following coculture. All data are shown as the means ± SD from three replicate cell cultures. P-values calculated by one-way ANOVA with Dunnett’s multiple comparison test (i). ns = non-significant.

Neutralizing anti-HSV-1 antibodies are usually directed against viral proteins that are displayed on the virion envelope. The neutralizing activity of anti-gB and anti-gD antibodies was assessed against cell-free virus in a pre-entry assay (**Fig. 5c**). Antibodies were incubated with HSV-1 virus, and pre-incubated virus was subsequently added to Vero E6 cells for infection. The antibody concentration was chosen to match IgG concentrations normally present in serum ^43^. Virus incubated with IgG control antibodies or treated with PBS served as negative controls. As a positive control, we included humanized antibody hu2c known to neutralize HSV-1 and to inhibit its cell-to-cell spread ^44^. Cells were fixed 4 h post infection and stained for HSV-1 immediate early protein ICP4 as a marker for infection. As expected, hu2c efficiently abrogated HSV-1 infection (**Fig. 5d**). Of the 10 human antibodies, three anti-gB antibodies and all five anti-gD antibodies efficiently prevented initiation of HSV-1 infection by cell-free virus (**Fig. 5d**).

HSV-1 escapes neutralizing antibodies by spreading directly to cells in close contact^45^. We employed a post-entry assay to assess the efficiency of anti-HSV-1 antibodies to inhibit viral spread by cell contact. Wildtype Vero E6 cells exposed to HSV-1 for 20 min were rinsed and overlayed with cellulose medium to restrict lateral viral spread to adjacent cells, allowing the formation of discrete plaques (**Fig. 5e**). Of the five anti-gB antibodies, one strongly inhibited intercellular virus spread, while two anti-gB antibodies showed a modest inhibitory effect. All anti-gD antibodies and the positive control antibody hu2c efficiently prevented intercellular spread of virus (**Fig. 5f; Supplementary Fig. S6**).

To test if antibodies that prevent intercellular spread of HSV-1 could also prevent he cell-to-cell spreading of misfolded Tau in our cell model, two antibodies with highest antiviral activity, one antibody against anti-gD (142-278) and one anti-gB antibody (142-204), were assessed for their ability to inhibit the intercellular dissemination of Tau aggregates. HEK Tau-GFP^AD^ donor cells were preincubated with human antibodies, hu2c, isotype control antibody or PBS alone (**Fig. 5g**). Importantly, control antibody hu2c and both human anti-HSV-1 antibodies efficiently inhibited intercellular Tau misfolding (**Fig. 5h**). The inhibitory effect on Tau transmission was also observed in cocultures with human primary astrocytes (**Fig. 5i, j**). We conclude that selected CSF-derived antibodies against HSV-1 gB and gD glycoproteins not only efficiently neutralize virions and block cell-to-cell spread of virus, but also abrogate intercellular dissemination of Tau aggregates.

## Discussion

HSV-1 is one of the most prevalent human viruses worldwide. Infections are usually acquired in early childhood and persist for life. There is increasing evidence to suggest that HSV-1 infection plays a role in the development of dementia. HSV-1 infection of neurons can directly trigger pathological events, such as increased production and accumulation of amyloid-β, as well as phosphorylation of Tau ^46, 47^. Recent findings suggest that phosphorylated Tau might even play a role in host defense against HSV-1 infection by binding to HSV-1 virion components and thereby neutralizing viral infectivity^19^. Recurrent reactivation of the latent HSV-1 in the CNS can also indirectly induce chronic inflammation, which is a major trigger of AD ^48^. Our data suggest a novel avenue by which latent HSV-1 in the CNS could directly contribute to proteopathic seed spreading, a phenomenon that is believed to underly the progressive spreading of protein misfolding throughout the brain ^49^.

HSV-1 establishes lifelong latency in sensory ganglia and CNS. During latency, transcription of HSV-1 lytic genes is drastically reduced. However, latency is more dynamic than expected, and expression of lytic genes in the CNS during latency has been documented in postmortem human samples ^22^ ^50^ and HSV-1-infected rodents ^51–55^. The lytic (active) HSV-1 replication process is divided into three basic transcriptional stages: immediate-early, early, and late. Expression of genes from all stages was reported in experimental latent infection ^51^ and in brains of AD patients and controls ^22^. The expression of lytic genes during latent CNS infection could theoretically also affect the spreading of proteopathic seeds throughout the brain by mediating interactions of viral glycoproteins with receptors on adjacent cells ^32, 33^. We here show that expression of the core HSV-1 fusion complex by donor cells harbouring protein aggregates is sufficient for the direct cell-to-cell spreading of protein misfolding. Expression of viral glycoproteins by donor cells increased total numbers of secreted EVs, in line with the reported increased EV secretion following HSV-1 infection ^56^. However, neither EVs nor CM of donor cells was effective at transmitting the aggregation phenotype to recipient cells, arguing that proteopathic seed spreading mainly involves close cell contacts. A recent study demonstrated that HSV-1 infection resulted in the EV-mediated release and subsequent uptake of phosphorylated Tau by primary neurons. ^57^. Spreading of phosphorylated Tau from HSV-1 infected to uninfected neurons in close proximity has also been shown in another study^19^. However, it is unknown if the transmitted Tau species were also seeding competent. As cell-to-cell transmission presents the predominant route of viral dissemination in vitro and in vivo, direct cell-to-cell contact likely also represents a plausible route for phosphorylated and aggregate Tau ^24, 44^.

The fact that HSV-1 glycoproteins could accelerate the intercellular dissemination of two independent proteopathic seeds demonstrates that this effect is cargo-independent. It is tempting to speculate that expression of HSV-1 lytic genes during latency in the CNS contributes to the prion-like spread of diverse proteopathic seeds associated with neurodegenerative diseases. Here, we used the NM prion-like domain of *S. cerevisiae* Sup35 as a model protein, which shares compositional similarity with domains in approximately 1% of mammalian proteins ^34^. Many of them, such as FUS or TDP-43, form disease-associated aggregates in neurodegenerative diseases, suggesting that HSV-1 glycoprotein expression could also affect spreading of those protein aggregates.

Glycoproteins gD and gB represent key components of the viral entry machinery and elicit potent HSV-1-neutralizing antibodies. We isolated B cells from CSF of HSV-1 encephalitis patients and identified several neutralizing antibodies against highly immunogenic viral envelope proteins^58^. We report cloning of CSF-derived human antibodies against gB and gD that potently neutralize HSV-1 virus and also efficiently block cell-to-cell transmission of proteopathic seeds. Interestingly, HSV-specific antibodies are not confined to encephalitis patients, but are prevalent in human CSF. Neutralizing anti-HSV-1 antibodies have already been found in the majority of CSF samples from individuals aged 5-9 years with no suspicion of HSV encephalitis, and these increase to over 90% in individuals aged 50 and over ^42^. It has been suggested that these CSF-derived antibodies originate from serum and not from intrathecal B cells ^42^. A recent study reported higher levels of anti-HSV-1 antibodies in the CSF compared to antibody levels in the serum of AD patients but not healthy controls ^59^. High CSF antibody levels were also positively correlated with p-Tau levels in the CSF ^59^. Importantly, neutralizing antibodies produced in response to infection usually fail to inhibit direct cell-to-cell infection ^60^. A large plasma screen of seropositive individuals recently demonstrated that only 5% of carriers mount a robust antibody response capable of inhibiting cell-to-cell spread of the virus ^61^. High titres of antibodies that can prevent direct cell-to-cell spread of HSV-1 have been reported to correlate with reduced recurrence of HSV-1^61^. Wether anti-HSV-1 antibodies in AD patients can inhibit cell-to-cell spread of virus and thus also likely the spreading of pathogenic protein aggregates remains to be established.

Since vaccines or antiherpetic drugs that can prevent the establishment of latency, eradicate the virus or prevent its reactivation are lacking ^62^, therapeutic antibodies that target viral glycoproteins represent an alternative strategy to combat HSV-1 morbidity and mortality^63^. To date, several antibodies have been reported to have antiviral efficacy against HSV-1 infection, few of which inhibit cell-to-cell spread of HSV-1 ^44, 60, 64, 65^. None of these antibodies have been tested for their efficiency in inhibiting intercellular proteopathic seed spreading. We propose that the here presented anti-HSV-1 antibodies could be used as a novel antiviral immunotherapy for the treatment of tauopathies.

## Methods

### HSV-1 encephalitis patients

Informed consent was obtained for all primary samples from the two patients with acute HSV-1 encephalitis, and the analyses were approved by the Charité - Universitätsmedizin Berlin’s Institutional Review Board (study protocol EA1/258/18).

### Antibodies

The following antibodies were used: Rabbit anti-gB (Leege, Klupp, unpublished), mouse anti-gD (sc-21719, Santa Cruz); rabbit anti-gH ^66^, rat anti-Sup35 M 4A5 (MABF3172, Merck); rabbit anti-GFP (Abcam); rat anti-HA 3F10 (11867423001, Roche); mouse anti-GAPDH 6C5 (ab8245, Abcam); rabbit anti-Tau (ab64193, Abcam); rabbit anti-V5 D3H8Q (13202, Cell Signaling); rabbit anti-Flotillin-1 EPR6041 (ab133497, Abcam); mouse anti-Hsp70/72 (C92F3A-5, Enzo); mouse anti-ICP4 (sc-69809, Santa Cruz). Human mOG.53 served as isotype control ^67^. Humanized anti-HSV-1 antibody hu2c ^44, 68^ detects HSV-1 and HSV-2 gB protein.

### Production of recombinant human anti-HSV-1 antibodies from cerebrospinal fluid

Patient-derived monoclonal antibodies were generated from cerebrospinal fluid (CSF) B cells using an established single-cell cloning strategy adapted from previously published methods ^69, 70^. Briefly, CSF was obtained from two patients diagnosed with acute HSV-1 encephalitis. Single B cells were isolated from CSF by fluorescence-activated cell sorting into 96-well PCR plates using the following gating strategy: lymphocytes (CD3⁻, CD14⁻, CD16⁻, DAPI⁻) were sorted into plates containing hypotonic lysis buffer and classified as antibody-secreting cells (ASCs; CD138⁺), memory B cells (MBCs; CD20⁺ CD27⁺), or non-memory B cells (NMBCs; CD20⁺ CD27⁻). The following antibodies were used for cell surface staining: anti-CD3-FITC (1:25; Miltenyi Biotec; #130-098-162), anti-CD14-FITC (1:25; Miltenyi Biotec; #130-098-063), anti-CD16-FITC (1:25; Miltenyi Biotec; #130-098-099), anti-CD20-PerCP-Vio700 (1:50; Miltenyi Biotec; #130-100-435), anti-CD27-APC-Vio770 (1:12.5; Miltenyi Biotec; #130-098-605), and anti-CD138-PE (1:50; Miltenyi Biotec; #130-098-122). The mRNA from individual B cells was reverse-transcribed to cDNA targeting the variable regions of immunoglobulin heavy and light chains. Subsequently, the variable regions of the antibodies were amplified by PCR. Sequence-confirmed variable domains were cloned into mammalian expression vectors containing human IgG constant domains. HEK293T cells were transiently transfected with plasmid pairs encoding matched heavy and light chains. Recombinant human antibodies were harvested from cell culture supernatants and purified using Protein G Sepharose beads.

### Plasmids

Generation of expression plasmids coding for the HSV-1 fusion machinery has been described previously ^66^. For detection of gL, a plasmid was generated in which the gL protein is tagged at the C-terminus with the V5 epitope tag GKPIPNPLLGLDST. Plasmid pMD2.VSV-G was from the Didier Trono lab. pHCMV was used as empty control vector. Plasmids expressing HVEM and Nectin-1 were kindly provided by Benjamin Vollmer and Kay Grünewald (CSSB, Hamburg, Germany) to the Mettenleiter lab.

### Cell lines

HEK NM-HA^agg^, HEK NM-GFP^sol^, HEK Tau-GFP^AD^ and HEK Tau-FR^sol^ cells have been described previously ^32^. HEK cells stably express a human Tau fragment that encompasses the four-repeat domain (4R) of human Tau (amino acids 243 to 375), which harbours P301L and V337M mutations. This fragment is fused to either GFP or FR (Evrogen), via an 18-amino acid flexible linker (EFCSRRYRGPGIHRSPTA), as previously described (hereafter referred to as ’Tau-GFP’ or ’Tau-FR’). HEK293T cells were cultured in Opti-MEM (Gibco), supplemented with glutamine and 10% (v/v) fetal bovine serum (FBS) (PAN-Biotech GmbH), as well as antibiotics. CHO-K1 cells were cultured in equal volumes of MEM (Hanks’ salts; Thermo Fisher Scientific) and MEM (Earle’s salts; Thermo Fisher Scientific), supplemented with 2 mM L-glutamine (Gibco), NaHCO₃ adjusted to 1.25 g/L, non-essential amino acids (NEAA; Gibco), sodium pyruvate (120 mg/L; Gibco) and 10% FBS at pH 7.2. Vero E6 cells were purchased from CLS (Cell Lines Service) and cultivated as recommended. Human primary astrocytes (ScienCell, #1800) were cultivated in accordance with the supplier’s recommendations. All cells were incubated at 5% CO₂ and 37 °C. The Vi-CELL XR Cell Viability Analyzer (Beckman Coulter) was used to determine the total numbers of viable cells. Cells were transfected using TransIT-2020/X2 (Mirus) or Lipofectamine 2000 (Invitrogen) reagents according to the manufacturer’s instructions.

### Production of recombinant NM

Competent *E. coli* BL21 (DE3) cells were transformed with a pET vector coding for the *Saccharomyces cerevisiae* Sup35 NM domain with a C-terminal His-tag under the control of the T7 promoter, at a concentration of 100 ng. The *E. coli* were transferred to 250 ml of LB medium containing 100 µg/ml ampicillin. NM-His expression was induced with 1 mM IPTG for 3 h at 37 °C with shaking at 180 rpm (Minitron shaker incubator, Infors HT). The pellets were pooled and lysed in buffer A (8 M urea, 20 mM imidazole in phosphate buffer) for 1 h at room temperature (RT). This was followed by sonication for 3 x 10 s at 50% intensity. The cell debris was pelleted for 20 min at 10,000 g before the supernatant was sterile-filtered. NM-His was purified from the supernatant via IMAC and HisTrap HP His tag protein purification columns (GE Healthcare) using the ÄKTA pure protein purification system (GE Healthcare). The column was rinsed with 75 ml of buffer A, after which NM-His was eluted using a linear imidazole gradient ranging from 10 mM to 250 mM (2–50% buffer B; 8 M urea and 500 mM imidazole in phosphate buffer). The pooled NM-His-containing fractions were concentrated using Vivaspin 20 concentrator columns (Sartorius, Germany) with a molecular cut-off of 10,000 Da. The protein was desalted using a 5 ml HiTrap Desalting Column (GE Healthcare) in sterile PBS and stored at -80 °C until further use.

### Production and transduction with lentiviral particles

Lentivirus was produced by transfecting HEK293T/17 cells with plasmids pRSV-Rev, pMD2.VSV-G, pMDl.g/pRRE, and pRRl.sin.PPT.hCMV.Wpre ^71^ containing NM-GFP or Tau-GFP/FR using Lipofectamine 2000 transfection reagent. The viral supernatant was harvested and concentrated using polyethylene glycol according to published protocols. The virus coding for NM-GFP was used to transduce CHO-K1 cells. A single cell clone was isolated that expressed NM-GFP uniformly. The virus coding for Tau-FR was used to transduce primary human astrocytes.

### NM aggregate induction in CHO NM-GFP^sol^ by recombinant Sup35 NM fibrils

To examine the capability of CHO NM-GFP^sol^ cells to form aggregates under the exposure of proteopathic seeds, 3 x 10⁴ CHO NM-GFP^sol^ cells per well were seeded in a 384-well plate and incubated in the presence of 25 µM of recombinant Sup35 NM fibrils. The next day, cells were fixed with 2% paraformaldehyde (PFA) and the nuclei were counterstained with 4 µg/ml Hoechst 33342 (Molecular Probes). The cells were imaged using a 20x objective with an automated confocal microscope (CellVoyager CV8000, Yokogawa Inc.). Maximum intensity projections were generated from Z-stacks. Images from 9 fields per well were taken. At least 9,000 cells per well and three technical replicates per treatment were analysed.

### Aggregate induction by coculture

For coculture experiments, the donor and recipient cells were cultured at a ratio of 5:1 on CellCarrier-96 or 384 black microplates (PerkinElmer). A total of 30,000 cells per well were plated. After 24 h, the cells were fixed with 2% PFA, and the nuclei were counterstained with 4 µg/ml Hoechst 33342 (Molecular Probes). The cells were imaged using a 20x or 40x objective with the automated confocal microscope CellVoyager CV8000 (Yokogawa Inc.). Maximum intensity projections were generated from Z-stacks. Images from at least 9 fields per well were taken. At least 3,000 cells per well and at least three technical replicates per treatment were analysed.

To culture HEK Tau-GFP^AD^ cells with astrocytes, human primary astrocytes were transduced with lentivirus encoding Tau-FR twice in 96-well plates within 5 d. HEK Tau-GFP^AD^ cells that were transiently expressing the HSV-1 viral fusion core, or that were mock-transfected, were seeded onto the astrocytes (at a donor-to-recipient ratio of 5:1). The average number of cells per well was 2.4 x 10⁴. The cells were cocultured for 24 h and were then fixed with 2% PFA. Nuclei were counterstained with Hoechst. The cells were imaged using a 20x objective. Maximum intensity projections were generated from Z-stacks. Images were taken from 9 fields per well using CellVoyager CV8000 (Yokogawa Inc.).

For testing anti-HSV-1 antibodies in the aggregate induction assay, the gBDHL- or mock-transfected donor HEK Tau-GFP^AD^ cells were pre-incubated 1 h at 37 °C with rotation at 20 rpm (Rotator RM-2M, neoLab) in the presence of 15, 1.5, or 0.15 µg/ml of recombinant human anti-gB / anti-gD antibody or humanized hu2c antibody ^44^. Incubation with isotype mGO.53 antibody at concentration of 15 µg/ml, PBS or medium was used for negative control samples. After incubation, the donor cells were combined with recipient cells (HEK Tau-FR^sol^ or astrocytes Tau-FR^sol^) at ratio of 5:1, and the coculture experiments were performed as above.

### Automated image analysis

An image analysis routine based on single cell segmentation and aggregate identification was developed using the CellVoyager Analysis support software (CV7000 Analysis Software; Version 3.5.1.18, Yokogawa Inc.). The total number of cells was determined based on the Hoechst signal. Recipients were identified based on their GFP/FR signal. Green or red aggregates were detected using morphological and intensity characteristics. The percentage of recipient cells with aggregated NM-GFP or Tau-FR was calculated by dividing the number of aggregate-positive cells by the total number of recipient cells set to 100%. The normalized percentage of recipient cells with aggregated Tau-FR for Fig. 5h was calculated as percentage of cells with red aggregates within a sample (e.g., incubated with an antibody) divided by percentage of cells with red aggregates in the sample incubated without an antibody; all values are presented in %.

### Immunofluorescence staining and confocal microscopy analysis

Confocal microscopy analysis was performed using either a Zeiss LSM 800 laser-scanning microscope with Airyscan (Carl Zeiss) or CellVoyager CV8000 (Yokogawa Inc.). The cells were fixed in 4% PFA, rinsed three times with PBS, and permeabilized either with 0.1% Triton X-100 (Zeiss LSM 800) or 0.5 % Triton X-100 (CellVoyager CV8000) for 10 min, or left untreated. The cells were blocked in 2% goat serum in PBS and incubated with primary antibodies in blocking solution for 2 h at RT or overnight (ON) at 4 °C. Cells were rinsed three times with PBS and subsequently incubated with either goat anti-mouse Alexa Fluor 647- (A-21235, Invitrogen), goat anti-rabbit Alexa Fluor 647- (A-11034, Invitrogen), or goat anti-human Alexa Fluor 647-conjugated secondary antibody (A-21445, Invitrogen). The nuclei were counterstained with 4 µg/ml Hoechst 33342 (Molecular Probes). The coverslips were mounted in Aqua-Poly/Mount (Polysciences). Confocal laser scanning microscopy was performed and images analysed using a Zeiss LSM 800 laser-scanning microscope with Airyscan (Carl Zeiss) and Zen2010 (ZenBlue, Zeiss). 384-well plates were scanned using a CellVoyager CV8000 (Yokogawa Inc.). Z-stacks were generated using optimal Z intervals (Nyquist criteria) and maximum intensity projections. ImageJ (Fiji) or CV7000 analysis software was used for image processing.

### Conditioned medium collection and EV isolation

Experiments with conditioned medium (CM) and EVs were performed using medium that contained EV-depleted FCS. FCS was centrifuged at 100,000 × g, 20 h, at 4 °C, to obtain EV-depleted FCS. Opti-MEM (Gibco) medium supplemented with the EV-depleted FCS and antibiotics was filtered through a 0.22 and a 0.1 µm filter (Millipore). To isolate CM and EV, 5 × 10^6^ HEK Tau-GFP^AD^ cells were seeded in 10 cm dishes in 10 ml of conventional Opti-MEM medium. The next day, the HEK Tau-GFP^AD^ cells were transfected with gBDHL or mock plasmids using Lipofectamine 2000. The medium was replaced with EV-depleted medium 5 h post-transfection. The conditioned medium was harvested 3 d post-transfection and centrifuged for 10 min at 300 × g at 4 °C (CM). For EV isolation, the CM was subjected to differential centrifugation (2000 × g, 20 min; 16,000 × g, 30 min; 100,000 × g, 1 h) using a SW45Ti rotor (Beckman Coulter). After centrifugation, the pellet was rinsed with PBS and centrifuged again using rotor SW55Ti (100,000 × g, 1 h). The EV pellet was resuspended in PBS and used in further experiments.

### Determination of size and number of EVs

The size and number of EVs were measured using a ZetaView PMX 110-SZ-488 nanoparticle tracking analyzer (Particle Metrix GmbH). Nanoparticle tracking analysis was conducted at 22 °C with adjustment of shutter and gain. At least three independent measurements were taken at 11 positions within the measurement cell for each sample. At least 1000 particles were tracked per measurement for all samples except for CM from mock-transfected samples where at least 350 particles per measurement were detected.

### HSV-1 infections of Vero E6 cells

HSV-1 strain lacking neurovirulence factor ICP34.5 and ICP47 blocking antigen presentation was used for infections (BSL1). Vero E6 cells, which are permissive, were exposed to HSV-1 at a multiplicity of 0.1 to 1 (MOI 1) and subsequently cultured for 24 h. The cells were fixed with 4% PFA and stained with recombinant human antibodies, HSV-1-specific antibodies, isotype mGO.53 or mouse anti-gD (sc-21719, Santa Cruz) to provide negative and positive controls, respectively. Goat anti-human Alexa Fluor 647-conjugated secondary antibody (A-21445, Invitrogen) was used to detect the binding of recombinant human antibodies.

### Virus pre-entry neutralization assay

The recombinant human monoclonal antibodies against gB or gD, an isotype mGO.53 control antibody, an anti-gB antibody hu2c ^44^ or PBS were incubated with HSV-1 ^67^ for 1 h at 37°C at a concentration of 15 μg/ml. HSV-1 (MOI 1) was added to Vero E6 cells grown in poly-L-lysine-coated 96-well plates, and the cells were fixed with 4% PFA 4 h post infection and stained for viral ICP4 (mouse anti-ICP4 antibody, sc-69809, Santa Cruz). The goat anti-mouse Alexa Fluor 488-conjugated secondary antibody (A-11029, Invitrogen) was used for ICP4 staining. The cells were imaged using the CellVoyager CV8000 (Yokogawa Inc.). Automated image analysis was performed using the CV7000 Analysis Software.

### Virus post-entry inhibition assay

Wild-type Vero E6 cells were plated at a density of 1.5 x 10⁴ cells per well in poly-L-lysine-coated 96-well plates. The following day, the cells were exposed to HSV-1 (MOI 0.01) for 20 min at 37 °C. The cells were rinsed with PBS and overlaid with 75 μg/ml antibodies in a medium containing 0.6% cellulose (#435244, Merck). The following day, the cells were fixed in 4% PFA for 1 h at RT. Cells were then rinsed two times with PBS, and the number of plaques per well was counted. Afterwards, the cells were rinsed once with PBS and then permeabilized with 0.5% Triton X-100 in PBS for 10 min at RT. Cells were then rinsed three times with PBS. The cells were then blocked in 2% goat serum in PBS for 1 h at RT. A primary mouse anti-gD antibody (Santa Cruz, sc-21719) was added overnight at 4 °C. The cells were washed three times and incubated with the goat anti-mouse Alexa Fluor 647-conjugated secondary antibodies (A-21235, Invitrogen) for 1 h at RT. The cells were rinsed three times again and stored at 4 °C before imaging.

### Western blotting

The Quick Start™ Bradford Protein Assay (Bio-Rad), the Fluostar Omega BMG plate reader (BMG Labtech) and the corresponding MARS data analysis software (BMG Labtech) were used to determine protein concentrations. Samples for SDS-PAGE were subjected to electrophoresis on NuPAGE® Novex® 4–12% Bis-Tris Protein Gels (Life Technologies). The proteins were transferred to a PVDF membrane (GE Healthcare) and detected by Western blot. Chemiluminescence (ECL, Thermo Fisher Scientific) was used to detect signals with primary antibodies and secondary goat anti-mouse IgG HRP (12-349, Dianova), goat anti-rabbit IgG HRP (12-348, Dianova) or goat anti-rat IgG HRP (AP136P, Dianova), in accordance with the manufacturer’s instructions, and the results were visualised using the Fusion FX imaging system (Vilber Lourmat).

### Statistical analysis

All statistical analyses were performed using Prism 6.0 (GraphPad Software v.7.0c). Two-sided unpaired Student’s *t-*test was used for 2-group comparison. We used one-way analysis of variance (ANOVA) with Dunnett’s post-hoc test for pairwise comparisons of multiple treatment groups with a single control group. P-values smaller than 0.05 were considered statistically significant. All experiments were repeated at least two times independently with similar results and performed in at least triplicate with similar results, except for anti gD antibody 142-313 in the post-entry assay (3 technical replicates). For quantitative analysis measurements were taken from distinct samples. Within our figures, the mean and the error bar representing the standard deviation (SD) are shown.

### Artificial intelligence tools

DeepL Write was used for text editing.

## Acknowledgements

Automated image analysis was performed in the laboratory automation technology facility, and confocal microscopy was performed in the light microcopy facility of the DZNE Bonn.

## Author Contributions

S.E.H., M.S., E.S.S., D.S., B.G.K., P.D., A. K. T.C.M., H.P. and I.M.V. contributed to the conception and design of the study; S.E.H., M.S., E.S.S., A.H., D.S, N.A., L. S., L.P., B.K., A.K. contributed to the acquisition and analysis of data; S.E.H., M.S., E.S.S., D.S., H.P. and I.M.V. contributed to drafting the text and preparing the figures.

