## Supplementary Figures for "The core herpes simplex-1 fusion complex drives cell-to-cell spreading of pathological Tau"

## S1

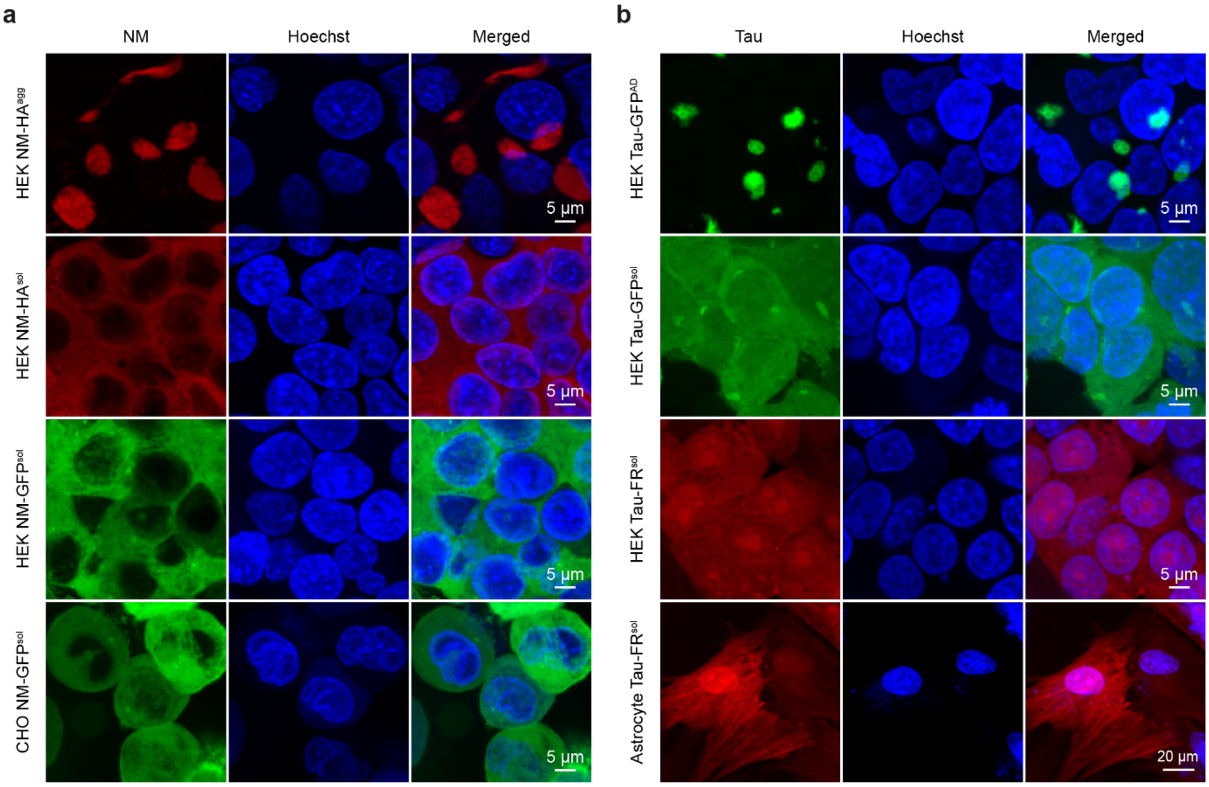


**Supplementary Figure S1. NM and Tau cell culture models. a.** HEK and CHO NM cell culture models. Shown are HEK and CHO cells expressing soluble NM-HA (NM-HA$^{sol}$) or soluble NM-GFP (NM-GFP$^{sol}$). HEK NM-HA$^{sol}$ cells had been exposed to recombinant NM fibrils to produce donors with NM aggregates. Following limiting dilution cloning, the donor clone HEK NM-HA$^{agg}$ was established that consistently produces NM-HA aggregates (NM-HA$^{agg}$) [1]. Nuclei were counterstained with Hoechst. **b.** Representative images of HEK Tau-GFP, HEK Tau-FusionRed (FR)$^{sol}$ and human primary astrocyte Tau-FR$^{sol}$ cell populations. HEK Tau-GFP$^{AD}$ cells stably express the four-repeat domain of human Tau with the familial P301L and V337M mutations fused to GFP. These donor cells were produced by exposing HEK Tau-GFP$^{sol}$ cells to AD brain homogenate, after which a single cell clone that persistently propagates Tau aggregates was isolated [1]. HEK Tau-FR$^{sol}$ cells express the Tau fragment fused to FR. The human primary astrocytes Tau-FR$^{sol}$ were generated by lentivirus transduction of astrocytes. Scale bars are indicated**.**

**S2**

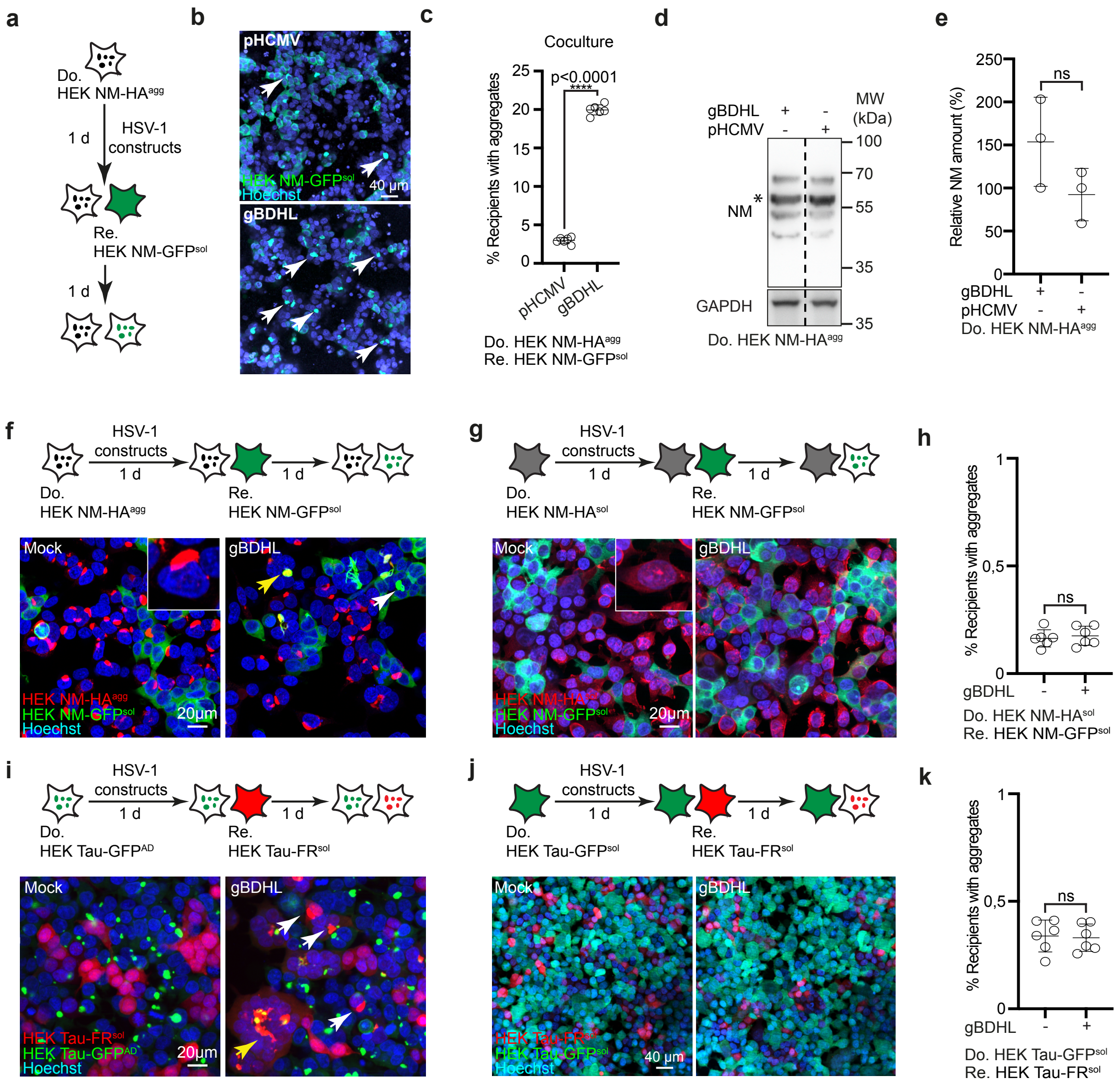


**Supplementary Figure S2. HSV-1 fusion machinery expressed by donor cells bearing aggregates increases aggregate induction in recipient cells. a.** Experimental design. HEK NM-HA$^{agg}$ donor cells were mock-transfected with pHCMV or transfected with all HSV-1 plasmids coding for gB, gD, gH and untagged gL. Cells were subsequently cocultured with recipient HEK NM-GFP$^{sol}$. **b.** Coculture of mock-transfected donors or donors transfected with plasmids coding for the HSV-1 fusion machinery with recipient cells. Arrowheads: NM-GFP aggregates. Nuclei were detected with Hoechst. Note that we have not stained for donor NM-HA in this experiment. **c.** Quantitative analysis of the percentage of recipient cells with induced NM-GFP aggregates. **d.** Western blot analysis of HEK NM-HA$^{agg}$ donor cells transiently transfected with HSV-1 fusion machinery or pHCMV (mock) control. Expression level of NM-

HA was detected using an antibody against the M domain of NM. The expected size of NM is indicated by a star. Additional lanes were excised for presentation purposes. **e.** For quantification of NM expression in HEK NM-HA$^{agg}$ donor cells, all antibody-positive bands were included. Mean expression levels in HEK donor cells transfected with mock control were set to 100 %. **f.** Cocultures of recipient HEK NM-GFP$^{sol}$ with donor HEK NM-HA$^{agg}$ cells transfected with either mock plasmid or all plasmids coding for the HSV-1 fusion machinery (for simplicity called gBDHL). NM-HA was detected by anti-HA antibodies. White arrowheads indicate aggregates. Yellow arrowhead denotes syncytia formation. **g.** As a control, recipient cells were cocultured with transfected HEK NM-HA$^{sol}$ cells. **h.** Quantitative analysis of the percentage of recipient cells with induced aggregates shown in (g). **i.** Coculture of HEK Tau-GFP$^{AD}$ donors transfected with mock plasmid or with plasmids coding for HSV-1 gBDHL with HEK Tau-FR$^{sol}$ recipient cells. White arrowheads indicate aggregates. Yellow arrowhead denotes syncytia formation. **j.** Control cocultures with transfected donors expressing soluble Tau-GFP. **k.** Percentage of recipient HEK Tau-FR$^{sol}$ cells with induced aggregates after coculture with HEK Tau-GFP$^{sol}$ donors transfected with HSV-1 gBDHL or mock plasmids. All data are shown as the means ± SD from six (c, h, k) replicate cell cultures. Three (c, h, k) independent experiments were carried out with similar results. P-values calculated by Student´s t-test (c, f, i). ns = non-significant.

**S3**

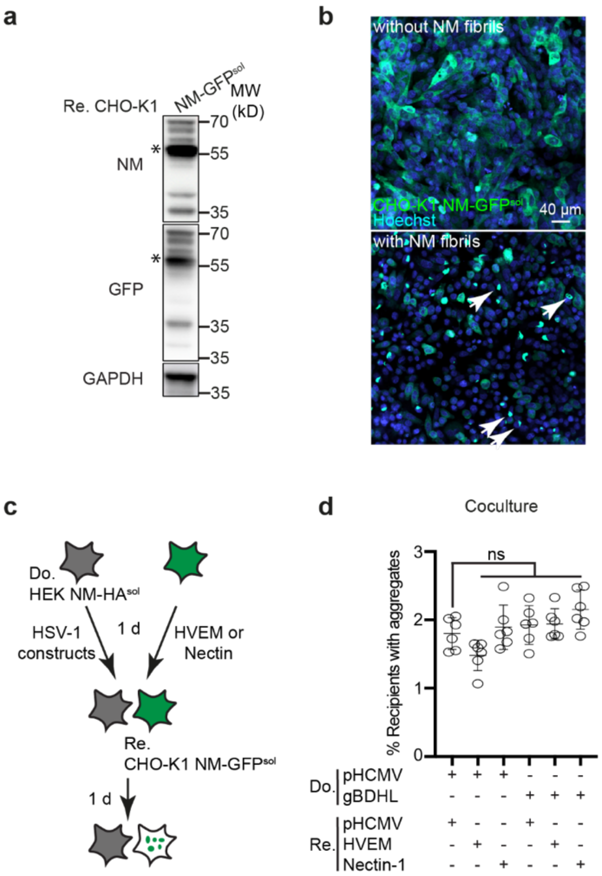


**Supplementary Figure S3. CHO -K1 cells expressing NM-GFP. a.** Western blot analysis of NM-GFP expression in CHO-K1 cells expressing soluble NM-GFP. NM-GFP was detected using an antibody against the M domain of NM and anti-GFP antibodies. The expected size is indicated by a star. Samples were loaded twice for GAPDH detection on a separate blot. **b.** CHO-K1 NM-GFP cell culture model. CHO-K1 cells expressing soluble NM-GFP (NM-GFP$^{sol}$, upper image) were exposed to 1 µM recombinant NM fibrils (monomer equivalent) and subsequently cultured for 24 h to confirm NM-GFP aggregate induction in these recipient cells (NM-GFP$^{agg}$, lower image). Arrowheads depict NM-GFP aggregates. **c.** Experimental workflow. As a control, donor HEK NM-HA$^{sol}$ cells expressing the HSV-1 fusion machinery were cocultured with recipient cells transiently transfected with plasmids coding for HSV-1 receptors HVEM, Nectin-1 or empty control vector pHCMV. **d.** Percentage of recipient HEK cells with induced aggregates. All data are shown as the means ± SD from six replicate cell cultures. Three independent experiments were carried out with similar results. P-values calculated by one-way ANOVA with Dunnett´s multiple comparisons test. ns = non-significant.

**S4**

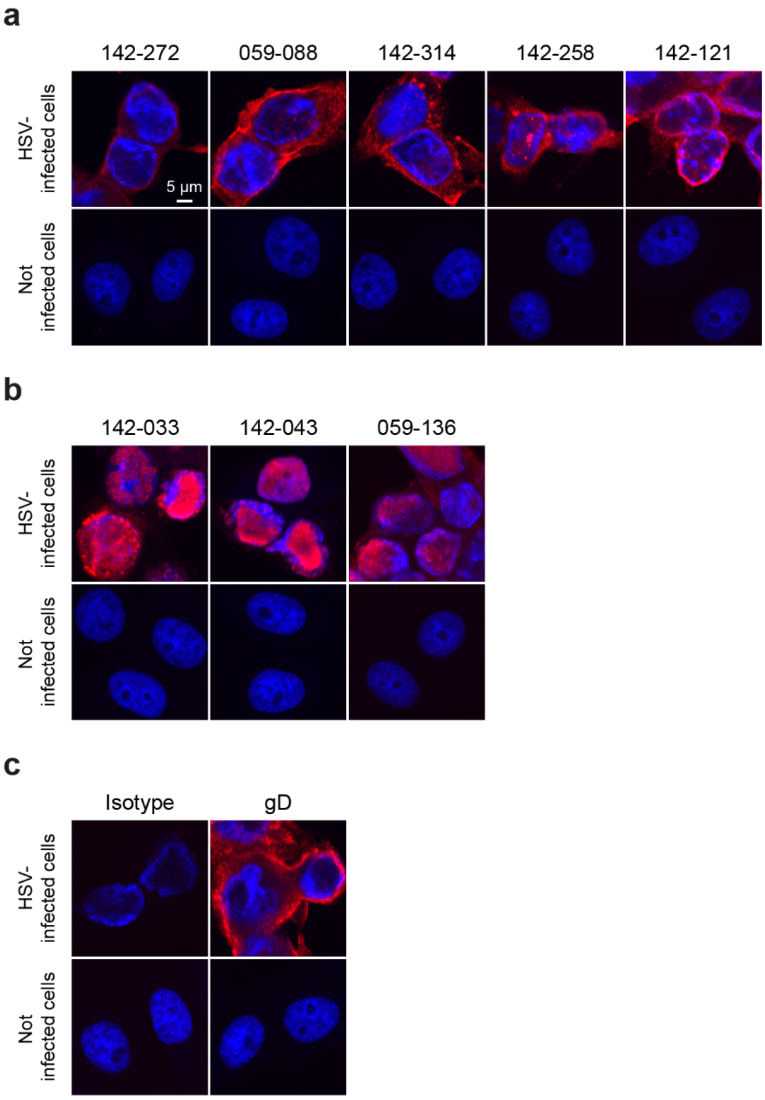


**Supplementary Figure S4. Detecting HSV-reactivity of patient CSF-derived antibodies. a-c.** Vero E6 cells were infected with HSV-1 (MOI 1) and fixed 24 h post infection. Permeabilized cells were stained with human monoclonal antibodies (**a, b**) or mouse anti-gD antibody (**c**). Nuclei were stained with Hoechst. Uninfected cells were stained with the same antibodies as controls. Of the 18 cloned antibodies, 3 detected nuclear targets (**b**).

S5

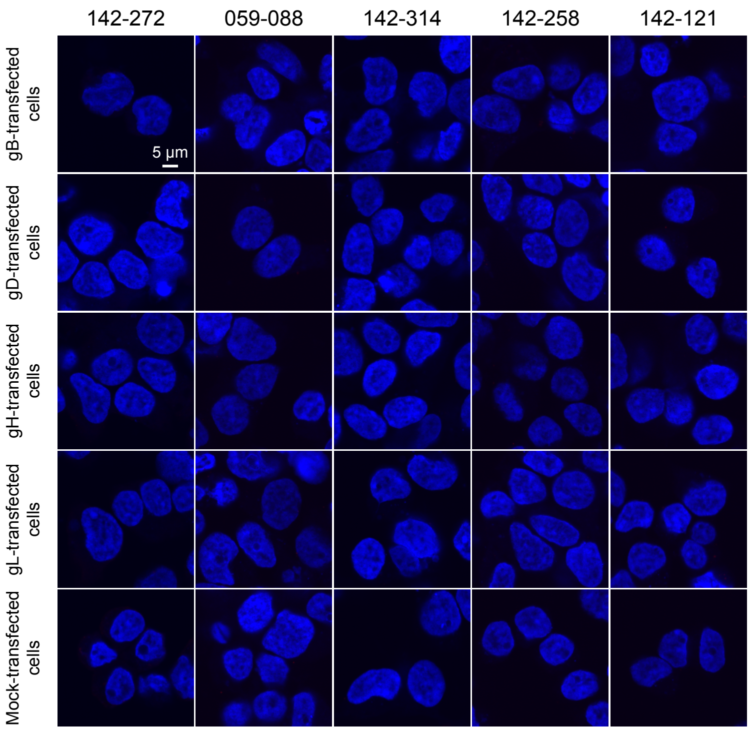


**Supplementary Figure S5. Antigen determination for patient CSF-derived antibodies.** HEK293T WT cells were mock-transfected (pHCMV control vector) or transfected with individual plasmids coding for HSV-1 gB, gD, gH, and gL and subsequently incubated with 5 human anti-HSV-1 antibodies to identify the antigen. Nuclei were stained with Hoechst. The here tested 5 antibodies did not detect gB, gD, gH, or gL.

**S6**

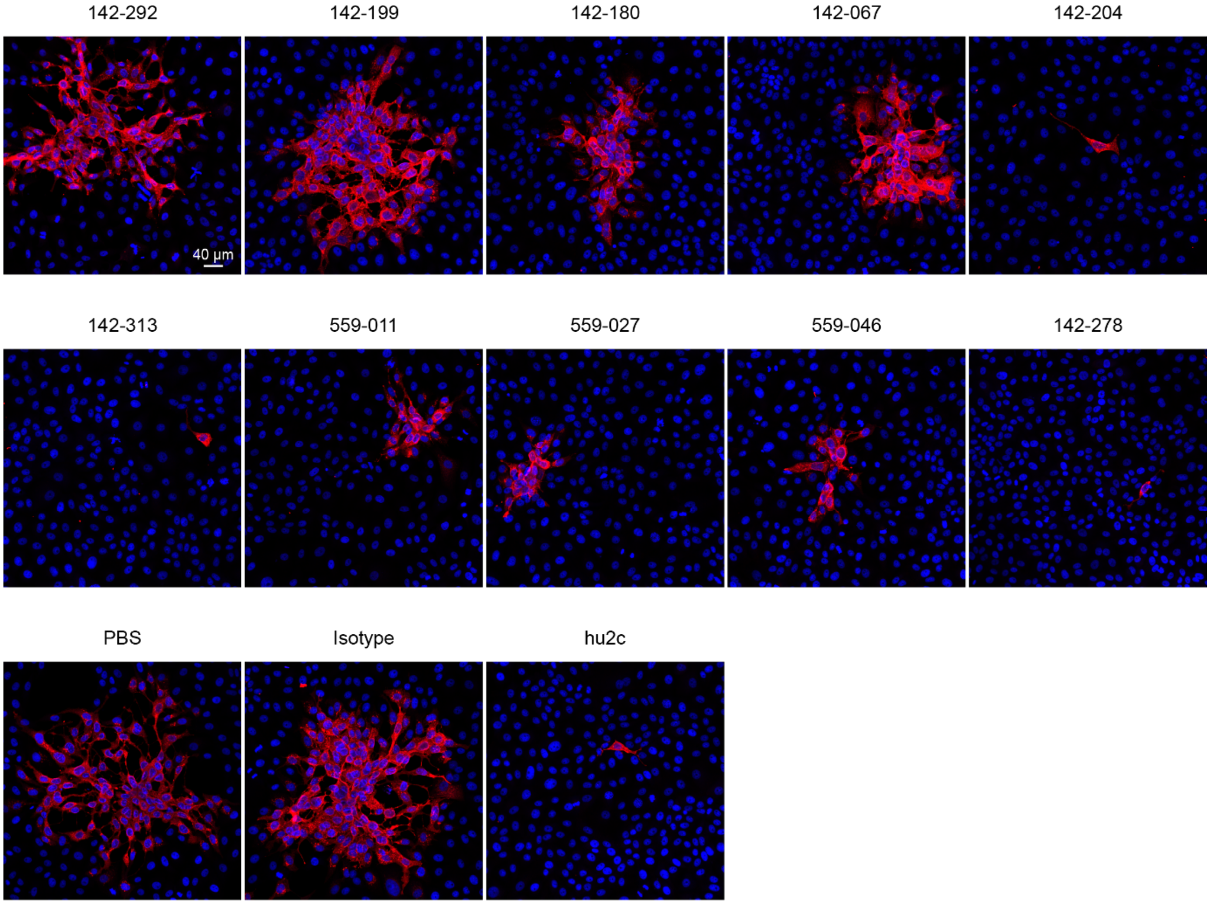


**Supplementary Figure S6. HSV-1 plaque formation in the post-entry assay.** Vero E6 cells were infected with HSV-1 (MOI 0.01). Cells were rinsed, and viral spreading was assessed 24 h after infection after incubating cells in the presence of 75 µg/ml anti-HSV-1 or control antibodies in 0.6% cellulose medium. Shown are plaques formed due to HSV-1 infection and stained with anti-HSV-1 gD antibody. Nuclei were stained with Hoechst.
